# Knockdown of CSNK2β suppresses MDA-MB231 cells growth, induces apoptosis, inhibits migration and invasion

**DOI:** 10.1101/2020.02.24.963769

**Authors:** Shibendra Kumar Lal Karna, Bilal Ahmad Lone, Faiz Ahmad, Nerina Shahi, Yuba Raj Pokharel

## Abstract

Breast cancer is most common cancer among women worldwide and among different types of breast cancer treatment of triple-negative breast cancer is major challenge, thus identification of specific drivers is required for targeted therapies of this malignancy. The aim of the present study is to elucidate the effects of silencing of CSNK2β gene by small interfering RNA (siRNA) on proliferation, cell cycle and apoptosis in breast carcinoma MDA MB-231 cells. Silencing of CSNK2β in MDA-MB-231(a triple negative cell line) cells resulted in decreased cell viability and colony formation. Cell cycle analysis showed that silencing of CSNK2β arrested MDA MB-231 cells in G2/M phase. We demonstrated that silencing of CSNK2β promoted nuclear condensation and augmented intracellular ROS production. Furthermore, Silencing of CSNK2β in MDA-MB 231 cells modulated the apoptotic machinery- BAX, Bcl-xL and caspase 3; autophagy machinary-Beclin-1 and LC3-1; and inhibited the vital markers (p-ERK, c-Myc, NF-κB, E2F1, PCNA, p38-α) associated with cell proliferation and DNA replication pathways. In addition, Knocking down of CSNK2 β also affected the migration potential of MDA-MB231 as observed in the wound healing and transwell migration assays. Together, our study suggests that CSNK2β silencing may offer future therapeutic target in triple negative breast cancer.

## Introduction

Casein Kinase 2(CSNK2), a highly conserved, multifunctional serine/threonine protein kinase, is critically important for the regulation of different processes in eukaryotes, such as proliferation, differentiation, and apoptosis^1^. CSNK2 is ubiquitously expressed in all tissues but its level is elevated in tumor tissues including prostate^2^, mammary gland^3^, head and neck^4^, lungs^5^ and kidney^6^. CSNK2 possess a heterotetrameric conformation with two catalytic and two regulatory subunits^7^. CSNK2β supports the structure of the tetrameric complex, augments catalytic activity and stability of CSNK2 and can also function dependently with other catalytic subunits^7^.

In mammalian cells, CSNK2β is phosphorylated at Ser 209 at its autophosphorylation site and at Ser 53B in a cell cycle-dependent manner^8^. CSNK2β is responsible for recruitment of CSNK2 substrates or potential regulators such as Nopp140, p53, Fas-associated factor-1 (FAF-1), topoisomerase II, CD5 and fibroblast growth factor-2 (FGF-2)^9–12^. Ectopic expression of CSNK2β in mouse 3T3-L1 adipocytes and in CHO cells increased proliferation^13^. The proliferative effects of CSNK2β vary in different cell lines. Deletion of gene encoding CSNK2β in mice leads to a failure in development^14^. In our previous work we have shown that CSNK2β ranks among top 10 proteins which increases likely hood of contributing oncogenic activity to its bait protein-Pin1^15^, a cis trans isomerase, and highly expressed in many cancer^16^. In the same work it has been found that siRNA knockdown of CSNK2β potently inhibited the oncogene Pin1, that supports the notion that CSNK2β is important for cancer pathogenesis. In a patient sample based study, specimens from 50 cholangiocarcinoma patients were evaluated for the expression of CSNK2β and XIAP by immunohistochemical staining where both showed strong correlation with tumor progression and CSNK2β was found significantly associated with TNM stage (P=0.036)^17^. In recent study, it has been revealed that traditional Chinese medicine huaier improves the survival rate of breast cancer patients by modulating the linc00339/mir-4656/CSNK2β pathways^18^. Together these reports suggest the significance of CSNK2β in cancer progression. To date, wealthy information is available on CK2α subunit and its signalling significance but very less is known about CSNK2β and its role as a signalling molecule particularly in context with the cancer cell’s proliferation and migration.

To understand its physiological importance in the regulation of multiple candidate target proteins, we focused our study on the role of CSNK2β in the tumorigenesis of human breast cancer (MDA-MB-231 cell) in vitro. In the present study, we used the RNA interference strategy to knockdown the CSNK2β gene and study the gross oncogenic activity in *an in-vitro* cell-based system. We evaluated its proliferative, clonogenic, invasive, and apoptotic properties in MDA-MB-231 cells using siRNA. We found that CSNK2β regulates the cell proliferation by targeting NF-κB, Wnt, and MAPK pathway proteins. Our findings suggest that CSNK2β can be used as a novel target for breast cancer therapy.

## Materials and methods

### Reagents

Lipofectamine RNAiMAX, TRIzol, Propidium Iodide, RNase were purchased from Invitrogen Corp (Carlsbad, CA, USA). siRNA was obtained from Qiagen (Hilden, Germany). Cell culture reagents and flasks were purchased from HiMedia (France) and Corning Inc (Corning, New York, USA). SYBR Green was purchased from Bio-Rad (Hercules, California). Antibodies were obtained from Santa Cruz Biotechnology (Dallas, Texas, USA), Cloud-Clone Corp. (Houston, USA). Cell Titer-Glo reagent was purchased from Promega Corp (Madison, Wisconsin, USA).

### Cell culture

MDA-MB-231 cell was purchased from National Centre for Cell Science (Pune, India). The cells were cultured in L-15 medium supplemented with 10% FBS, penicillin (100 unit/ml) and streptomycin (100µg/ml). The cell culture was maintained at 37°C in humidified air containing 5% CO_2_.

### Transfection

Cells were cultured in 6 well plates one day before siRNA transfection. We used 25 nanomolar of each siRNA and made complex in Opti-MEM media. Similarly, the complex of Lipofectamine RNAiMAX (4 µl/each well) and Opti-MEM was made and incubated for 5 minutes at room temperature. After that, both the complexes were mixed in 1:1 proportion and incubated for 25 minutes at room temperature. Cells were treated with Opti-MEM-siRNA Lipofectamine complex and incubated at 37°C for 72 hours. The sequences used in siRNA and the target sequence for the genes in our study are mentioned below.

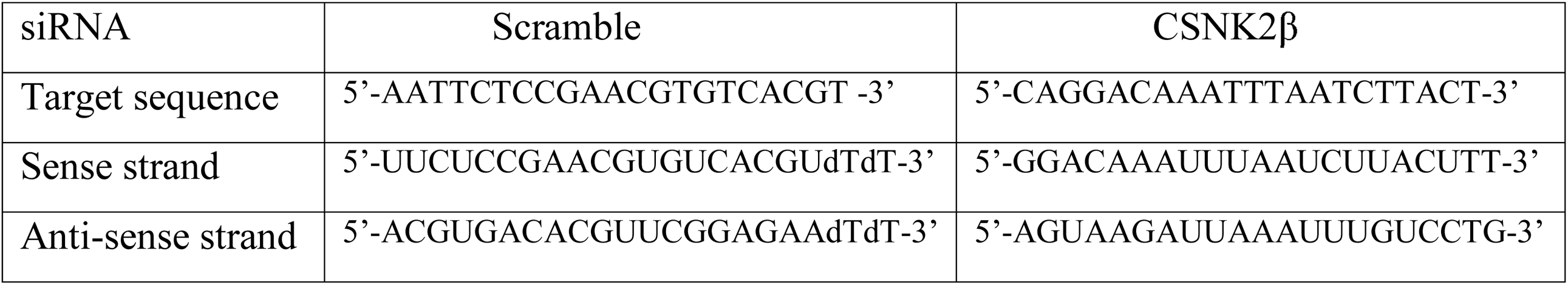

### Cell viability assay

Cell viability was assessed with CTG assay (Promega, Madison, WI). Briefly, MDA-MB-231 cells were seeded in 96 well white culture plate at a density of 3000 cells per 180 µl of medium per well with 20 µl of siRNA complexes for CSNK2β and incubated at 37°C, 5% CO_2_ for 24 hours. On the next day, the media containing the complex was changed with the fresh media and further incubated till 96 hours. The cells were treated in quadruplets with respective siRNA. The reagents were prepared according to manufacturer’s protocol. After incubation 100 µl of fresh media was added to each well followed by 100 µl of reagent and kept on a shaker for 2 minutes to induce the cell lysis. The plate was incubated for 10 minutes at room temperature to stabilize the luminescence signal. Luminescence was measured using microplate ELISA reader (Bio Tek, Winooski, Vermont, US).

### Colony formation assay

MDA-MB-231 cells were transfected with Scramble and CSNK2β siRNA and incubated for 48 hours. After the cells were trypsinized, collected and counted. 500 cells/well were taken from each Scramble and CSNK2β transfected samples and seeded in 6 wells plate. Every two days media was changed and the cells were allowed to grow for 3 weeks. Then the cells were washed with DPBS, fixed with 3.7% formaldehyde for 10 min and stained with 0.4% crystal violet. The cells washed DPBS for 2-3 times and allowed to dry. The colonies were counted using Image J software.

### Wound healing assay

MDA-MB-231 cells were plated in 6 well plates (4×10^5^ cells/well) and transfected with Scramble and CSNK2β siRNA as mentioned above. After the cells reach 90% confluence, a wound was made in the center of the plate by scrapping the cells monolayer with 10µl tip. The cells were washed with sterile DPBS to remove the detached cells. The width of wound area was measured at different time points using an inverted microscope. Densitometry analysis of wound area was carried by using Image J software

### Matrigel Invasion assay

Transwell chambers (corning) were used for invasion assay. MDA-MB-231cells were transfected with CSNK2β siRNA and post 48 h of transfection, cells (5 × 10^4^) were harvested and transferred to the upper chamber of transwell, pre coated with matrigel, in 200 µL of serum free L-15 media whereas lower chamber consisted of complete media serving as a chemoattractant for the cells. Following 24 h of incubation, cells in the upper surface of chambers were removed with cotton the cells invaded through matrigel at lower surface were fixed in 70% ethanol and stained with 0.4% crystal violet. Quantification of invaded cells was done by counting in six random fields.

### Cell cycle analysis

2.5×10^5^ cells were seeded in 6 well plates and transfection was done as mentioned before. The cell cycle analysis was performed 48 hours post-transfection using BD FACS Verse ™. The cells were pooled out and fixed with ice-cold 70% ethanol and kept overnight at 4°C. On the next day, the cells were washed with DPBS for two times. The cells were further digested with RNase A (50 U/ml) at 37°C for 1 hour and then stained with PI (20 μg/ml) at room temperature for 20 min. The samples were acquired by flow cytometer (FACS Verse) and analysed by ModFit software.

### Western blotting

MDA-MB-231 cells were cultured in 6 well plates. On the next day cells were transfected with Scramble and CSNK2β siRNA and incubated for 72 hours. After incubation, the cells were washed twice with ice-cold DPBS and then lysed with sodium dodecyl phosphate (SDS) lysis buffer. The samples were heated for 5 minutes at 100°c. The samples were run on SDS-PAGE, transferred to PVDF membrane (90 volts for 2 hour). The membranes were blocked using 5% skim milk in Tris buffer saline with 0.1% tween-20 (TBST) for 2 hours at room temperature. The blots were incubated with primary antibodies overnight at 4°c on a rocker followed by incubation with HRP conjugated secondary antibodies. The blots were developed using enhanced chemiluminescence (ECL, Biorad). β actin was developed as a loading control.

| S/NO. | ANTIBODY | MANUFACTURER | DILUTION |
| --- | --- | --- | --- |
| 1 | β-Integrin | Cloud clone corp. | 1:500 |
| 2 | E-Cadherin | Cloud clone corp. | 1:500 |
| 3 | Vimentin | Cloud clone corp. | 1:500 |
| 4 | MMP9 | Cloud clone corp. | 1:500 |
| 5 | β-Actin | Cloud clone corp. | 1:2000 |
| 6 | Bax | Santa Cruz Biotechnology Inc. | 1:500 |
| 7 | Bcl-xl | Santa Cruz Biotechnology Inc. | 1:500 |
| 8 | Caspase 3 | Santa Cruz Biotechnology Inc. | 1:500 |
| 9 | LC-3 | Cloud clone corp. | 1:500 |
| 10 | Beclin | Cloud clone corp. | 1:500 |
| 11 | CSNK2β | Santa Cruz Biotechnology Inc. | 1:1000 |
| 12 | PCNA | Cloud clone corp. | 1:1000 |
| 13 | c-myc | Santa Cruz Biotechnology Inc. | 1:1000 |
| 14 | NfκB | Santa Cruz Biotechnology Inc. | 1:1000 |
| 15 | E2F1 | Santa Cruz Biotechnology Inc. | 1:500 |
| 16 | p-erk | Santa Cruz Biotechnology Inc. | 1:500 |
| 17 | p-38 | Santa Cruz Biotechnology Inc. | 1:500 |

### RNA extraction, cDNA preparation, and real time PCR

Cells were transfected for 72 hours with the respective siRNAs and RNA was extracted using TRIzol lysis reagent using manufacturers protocol. 2µg of RNA was taken and pretreated with DNase followed by cDNA synthesis with oligoDT primers. The thermocycler condition was 65° c, 5 min, 42°c, 60 min and 72°c (final extension). Real time PCR was done with iTaq SYBR green mix (Biorad) using ABI 700,(Invitrogen). Primer sequences are given below

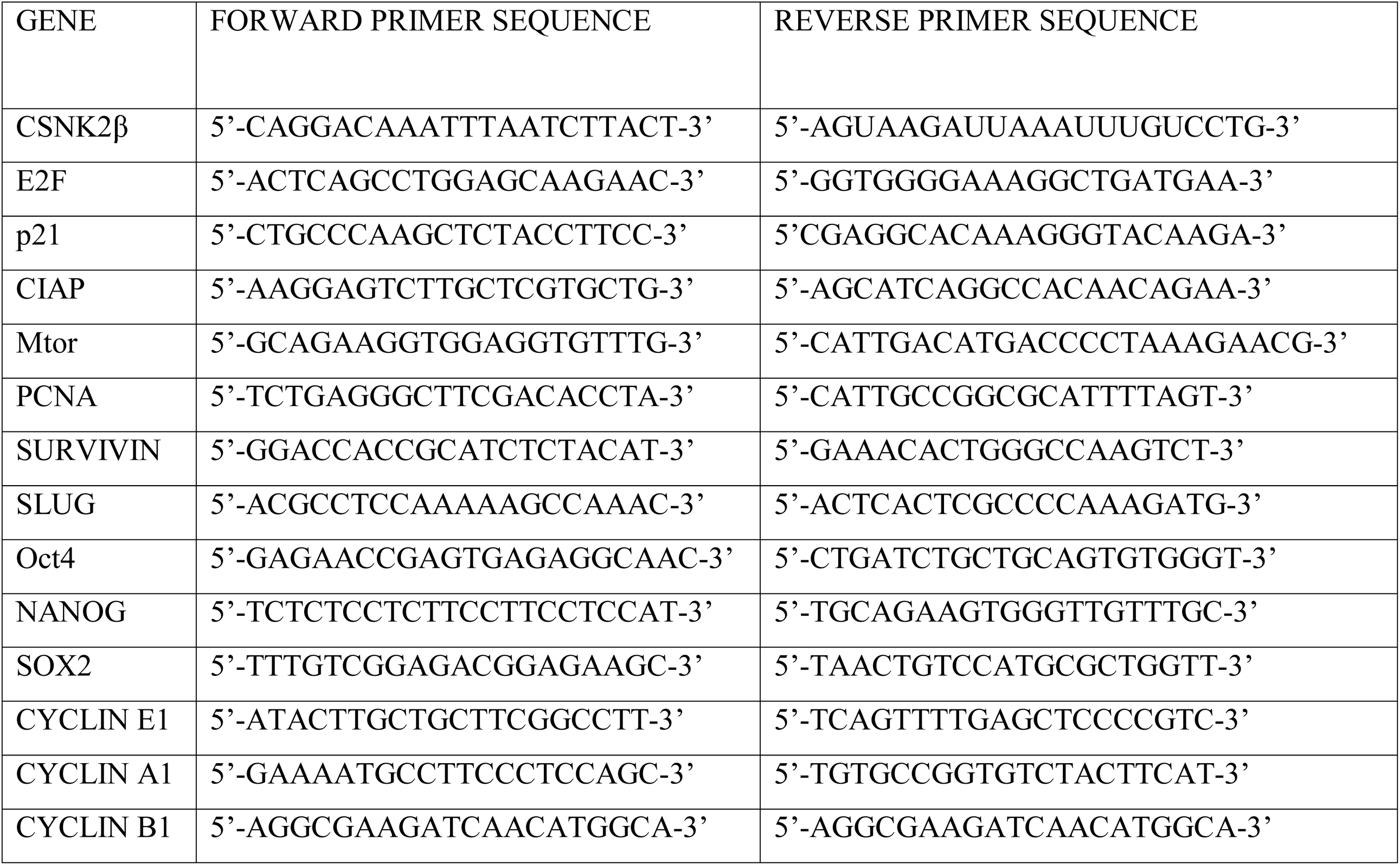

### Hoechst staining

1.5×10^5^ cells were seeded in 6 well plates. On the next day, the cells were transfected with Scramble and CSNK2β siRNA and incubated for 72 hours. Then the cells were stained with Hoechst 33342 for 15 minutes in dark. Cells were once washed with DPBS and the images were taken using Nikon fluorescent microscope.

### Apoptosis assay

Cell apoptosis assay was performed by flow cytometry using an Annexin V-FITC /propidium iodide(PI) apoptosis detection kit(BD). Cells were seeded in triplicates in 6-well plate at the density of 2×10^5^ cell per well. After 48 hours of transfection with Scramble and CSNK2β siRNA, the cells were harvested and counted and the Annexin V-FITC/PI assay was performed according to the manufacturer’s instruction. A total of 30,000 cells were analysed using FACSuite software. The percentage of AnnexinV+/PI+ cells in the total cell number was used as the apoptosis cells.

### Measurement of the intracellular ROS production

Intracellular ROS generation was assessed by fluoroprobe CM-H2DCFDA(Invitrogen). Approximately 1.5×10^5^ cells were cultured in 6 well plates and transfected with the Scramble and CSNK2β siRNA for 72 hours. The cells were treated with 10 µM CM-H 2 DCFDA for 30 min in the dark at 37°C. The wells were once washed with DPBS and the images were captured using a fluorescence microscope (Nikon Ti).

### Statistical Analysis

Data were represented as a mean ± standard deviation. The level of significance between two groups was calculated by t-test. P < 0.05 was considered statistically significant.

## Results

### Comparative analysis of CSNK2β expression in multiple cancer types and the correlation with overall survival

To understand the expression status of CSNK2β in different cancers, we used the datasets retrieved from cbioportal^19^ and compared the genomic alterations in multiple tumours. CSNK2β showed high degree of amplification in most cancer type followed by deletion and mutation (Figure 1A). Data so obtained from cbioportal shows that about 14% gene amplification of CSN2β in breast cancer. Next, we checked the status of CSNK2β in different cancer from GEPIA database^20^. Among 32 different cancers showed in the panel, expression is significantly higher in bladder urothelial carcinoma, breast invasive carcinoma and mesolthelioma (Figure 1B). Each dot represents expression of samples. Data for breast cancer lie in the third column. We also observed the distribution of CSNK2β gene expression between breast tumor tissue and adjacent tissue using MERAV (http://merav.wi.mit.edu/)^21^. As shown in (Figure 1C) the expression of CSNK2β is upregulated in breast tumor. Next, mRNA expression analysis of CSNK2β in a large cohort of datasets containing 3951 breast cancer patients with an online tool KM-Plotter (http://kmplot.com/analysis)^22^ showed that higher expression of CSNK2β in breast cancer patients is correlated with reduced overall survival (p<0.001, longrank) (Figure 1D)^23^. Together, these results suggest that CSNK2β is clinically important for cancer pathogenesis and correlates with patient outcome.

**Figure 1.**
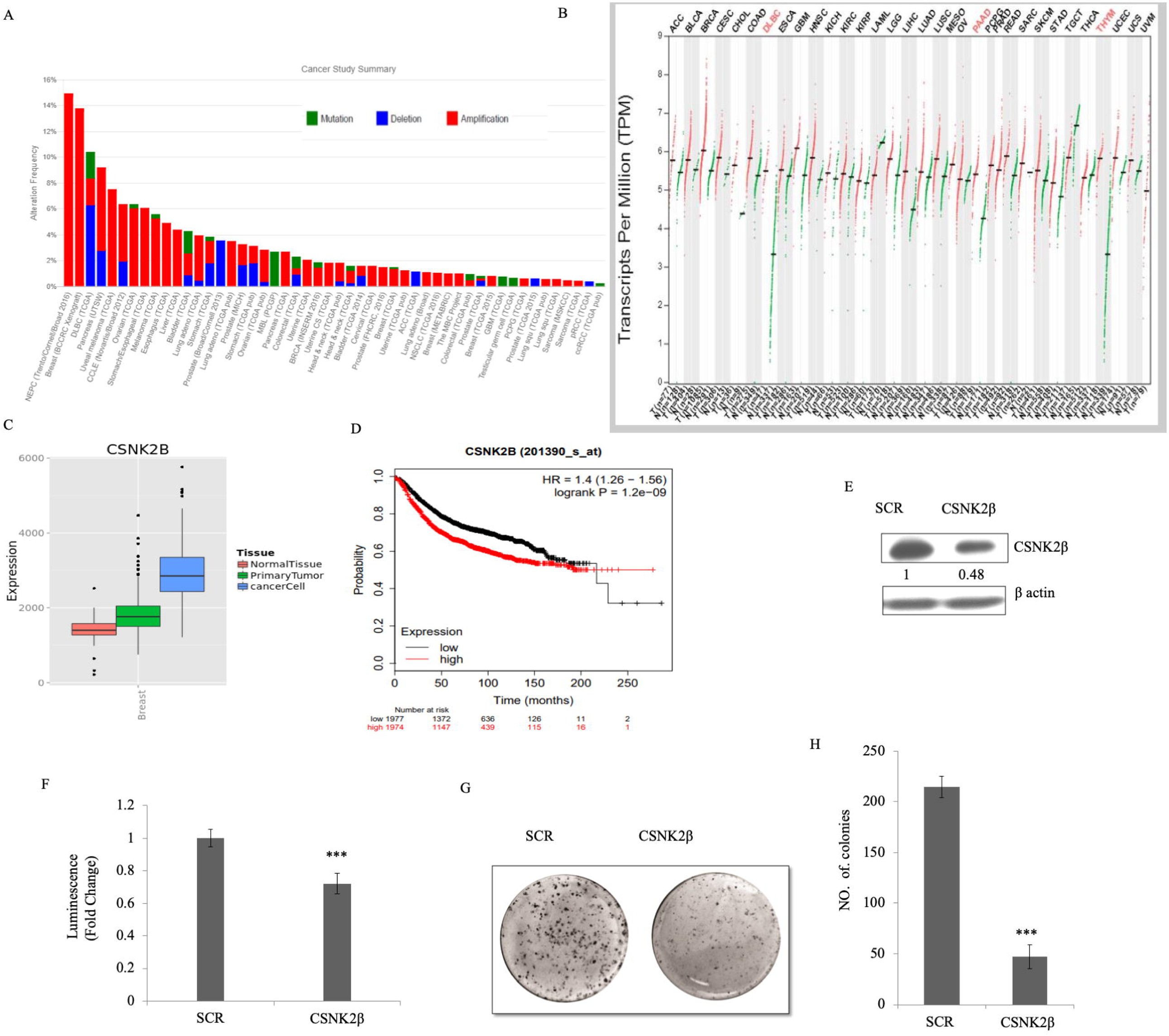
CSNK2β expression in multiple cancer types and its functional role in breast cancer cell proliferation. **(A)**. Mutational status of CSNK2β across human cancers based on cBioPortal data. Histogram representing the mutational landscape (mutation, amplification, and deep deletion) of CSNK2β in different types of cancer **(B)** The gene expression profile across all tumor samples and paired normal tissues. For each tumor (red), its matched normal(green) are given; T: tumor; N: normal; n: number. Y axis represents transcript per million in log scale whereas X axis represnts number of tumor and normal samples **(C)** Comparison of gene expression between breast tumor tissue and adjacent normal tissue using MERAV database. **(D**) Kaplan-Meier survival curve of the mRNA expression of CSNK2β for breast cancer patients(n=3951) using KM-plotter database. **(E and F)** Silencing of CSNK2β and evaluation of its expression by western blot followed by cell viability, assayed by CTG assay **(G and H)**. Knockdown of CSNK2β decreases the colony formation potential of MDA-MB-231 cells. Bar graphs depict the number of colonies in each sample. Data are represented as mean ± SD from triplicate samples, where *** denotes p < 0.001.

### Silencing of CSNK2β inhibited proliferation of MDA-MB-231cells

To study the knocking down effect of CSNK2β in vitro, we silenced the expression of CSNK2β using siRNA approach and evaluated the CSNK2β protein level (Figure 1E) which confirmed the siRNA efficiency. Next cell viability was measured 96 hours of post-transfection by using CTG substrate, the viability of CSNK2β silenced MDA-MB-231 cells was significantly lower as compared to scrambled siRNA (p< 0.001) (Figure 1F). These results suggested that CSNK2β play a crucial role in cell proliferation of MDA-MB-231 cells. Clonogenic assay which mimics the tumor formation *in vivo* was carried out to observe the long term survival of cells, which were transfected with CSNK2β +/-siRNAs. Silencing of CSNK2β significantly reduced the number of colonies as compared to Scramble siRNA transfected samples (p< 0.001) (Figure 1G & H). This result further confirmed the pivotal role of CSNK2β in the proliferation of MDA-MB-231 cells.

### Silencing of CSNK2β augmented ROS production, caused cell cycle arrest at G2/M and induced apoptosis in MDA-MB-231 cells *in vitro*

ROS possess double edge sword property both as, oncogenic-maintaining sustained and increased proliferation of cancer cells, as well as, tumour suppressor-leading to cell death, when an aberrant increase in intracellular ROS arises due to any kind of stress. It is a good idea to find oxidative stress modulators as an anti-cancer strategy. In our experiment, we found the increased production of ROS in CSNK2β siRNA transfected cells compared to scramble siRNA transfected cells (Figure 2A). This data suggested that CSNK2β play crucial role in attenuating the intracellular ROS production in MDA-MB-231 cells for its sustained proliferation.

**Figure 2.**
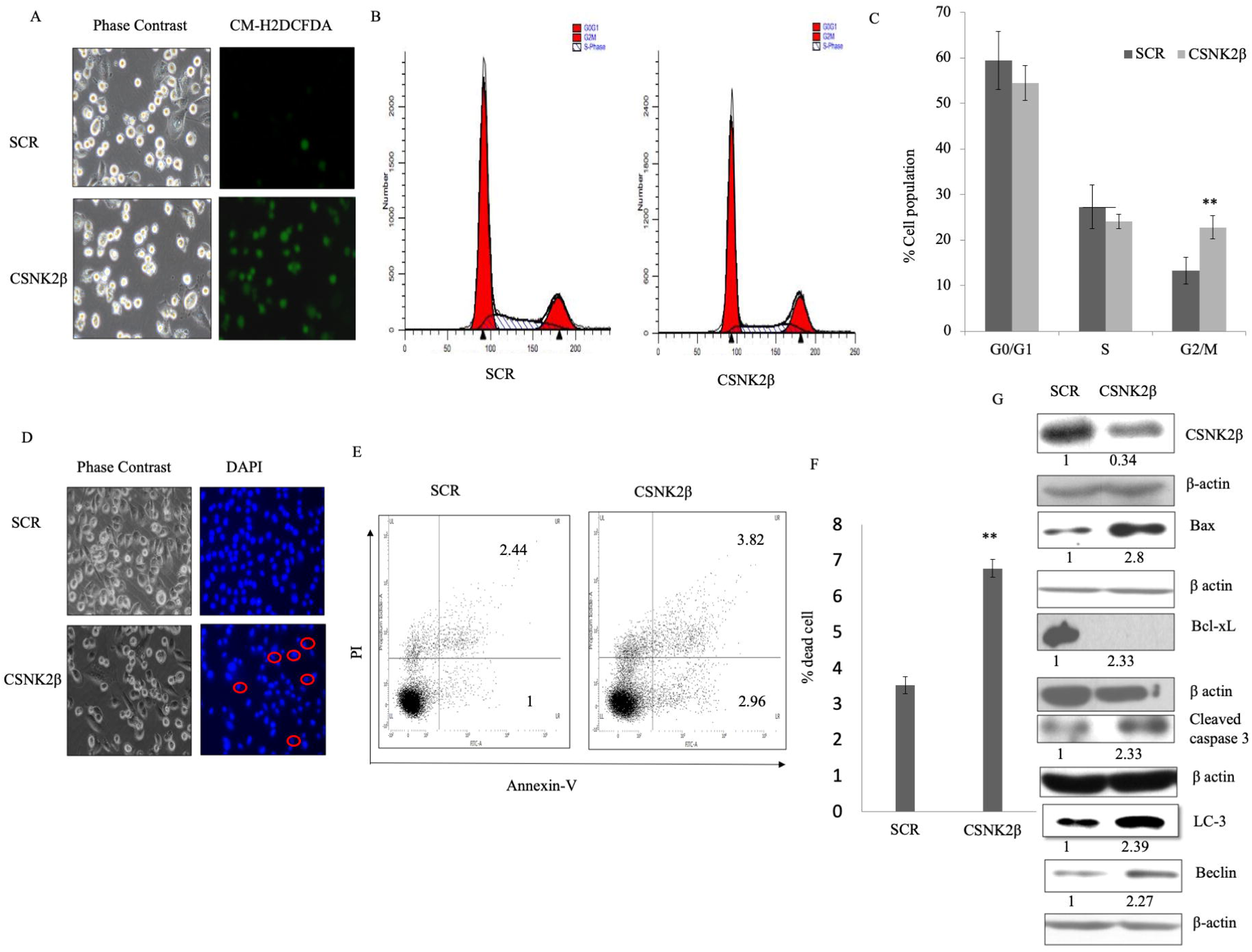
Silencing of CSNK2β induced apoptosis in MDA-MB-231 cells *in vitro*. **(A)** ROS detection-fluorescent microscopic images of MDA-MB-231 cells stained with CM-H2DCFDA, following knockdown of CSNK2β. **(B and C)** Histogram and bar graph represent the percentage of MDA-MB-231cells in different stages of cells following silencing of CSNK2β **(D)** Microscopic evaluation of cell morphology and Hoechst stained nuclei. Condensed nuclei of apoptotic cells are circled as characterised by smaller and fragmented appearance with respect to control. **(E and F)** Annexin V-FITC/PI double staining for apoptisis. Bar graph shows total apoptotic cells (early+late apoptosis).**(G)** Western blot analysis of apoptosis associated markers. Knockdown of CSNK2β inhibited Bcl-Xl whereas activated Ba, caspase 3, Beclin and LC-3. Results for Annexin V-FITC/PI double staining are representative of two independent experiments represented as a mean ± standard deviation, where **p < 0.01.

Next we checked the status of cell cycle phase following knockdown of CSNK2β. After 48 hours of transfection, the percentage of cells in the different phase of the cell cycle was analyzed by flow cytometry. The percentage of cells in G2/M phase was significantly increased in CSNK2β transfected samples (22.806%) than in scrambled siRNA transfected samples (13.266%)(p < 0.01). At the same time, the percentage of cells in G0/G1 and S phase was reduced (Figure 2 B & C). These results suggest that knockdown of cells with CSNK2β inhibits the progression of cells via blocking the cell cycle at G2/M phase.

The significant arrest of MDA-MB-231 cells in G2/M phase led us to check the effect of silencing CSNK2β on cell apoptosis. We found that silencing of CSNK2β caused change in cellular morphology of cells and condensation of nuclei-a prominent feature of cells undergoing apoptosis (Figure 2D). A quantitative estimation of apoptotic cells was carried out using FACS and the result of FACS analysis with double positive annexin V-FITC/PI showed 6.77% (early apoptotic+late apoptotic population) in CSNK2β siRNA sample as compared to 3.52% (early apoptotic+late apoptotic population) in scramble siRNA sample which comes out to be significant with p < 0.01 as shown in (Fig 2E & F). To unravel the cell death mechanism induced after the knockdown of CSNK2β, western blot analysis was performed to determine the level of proteins related to apoptosis. We found that, there was increased expression of BAX (pro-apoptotic), and a decrease in the expression level of Bcl-xL (anti-apoptotic), increased expression of cleaved caspase 3 (Figure 2 G). Altogether these results showed that silencing of CSNK2β drives cell death via apoptosis. Moreover, we also analyzed the expression of two vital autophagy markers; Beclin and LC3, and found both of the markers upregulated in CSNK2β siRNA transfected samples than Scramble siRNA (Figure 2G). These findings showed that targeting CSNK2β in MDA-MB-231 triggers the cells towards both apoptosis and autophagy cell death pathway

### Knockdown of CSNK2β inhibited migration and invasion of MDA-MB-231 cells

Next we questioned whether depletion of CSNK2β could alter the migration and invasion of MDA-MB-231 cells. In wound healing assay, migratory capacity was observe for each sample at different time points (0, 6 and 12h). After 12 hours, the area of a wound was significantly wide in CSNK2β transfected wells as compared with scramble suggesting the role of CSNK2β gene in the migration of MDA-MB-231 cells *in vitro*. (p < 0.001)(Figure 3A & B). Next we also checked the invasive potential of MDA-MB-231 following CSNK2β knockdown and found that cells transfected with siRNA CSNK2β were unable to invade the matrigel layer as efficient as the siRNA Scramble transfected cells were able to transverse across the matrigel layer, shown in the (Figure 3C & D). Some critical markers associated with cell migration were also checked and it was found that knockdown of CSNK2β, caused inhibition of MMP-9, Vimentin,Integrin and β-catenin whereas it upregulated E-cadherin (Figure 3E).

**Figure 3.**
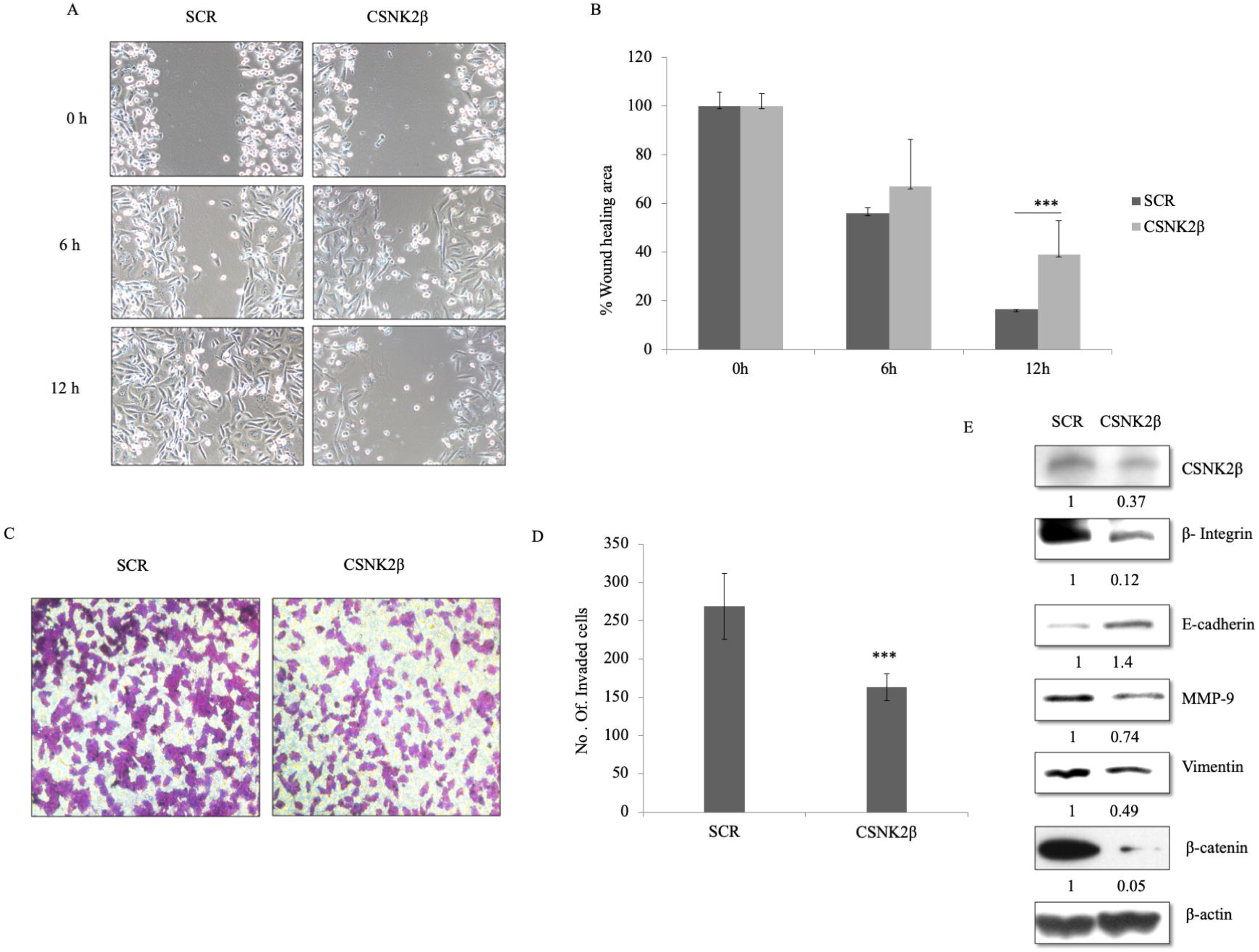
Silencing of CSNK2β inhibited migration and invasion of MDA-MB-231 cells. **(A and B)** Representative images of wound healing assay performed to evaluate the motility of cells after silencing CSNK2β. After transfection, a scratch was made on cells monolayer and was monitored with microscopy every 6 hours (0, 6 and12h). Bar graphs show normalized wound area, calculated using Image J software. **(C and D)** Representative images of invasion assay. Bar graphs depict number of invaded cells following knockdown of CSNK2β Data are represented as mean ± SD from triplicate samples, where **p <0.01 and ***p < 0.001. **(E)** Western blot analysis depicts protein expression of migration associated markers. Knockdown of CSNK2β up regulated E-cadherin whereas down regulated integrin,mmp-9 and β-catenin.

### Silencing of CSNK2β modulated key markers associated with cell proliferation, migration and cell stemness

To elucidate the role of CSNK2β in MDA-MB-231 cell’s proliferation, survival and apoptosis we examined the expression of some selected genes at mRNA and protein level related to these processes by real time PCR and western blotting respectively. We checked the expression of some genes, well known for their involvement in cancer progression, at protein level. Knockdown of CSNK2β led to down regulation of PCNA,E2F1,c-MYC,NF-Kb,p-Erk and p-38α as shown in (Figure 4A). Next we were interested to check the expression of few other critical genes at mRNA level. In 72 hours transfected samples we found that the mRNA level of p21, a cell cycle inhibitor, was increased up to about 2.7 fold, while the expressions of survivin (anti-apoptotic and metastatic marker), Nanog, SLUG, OCT4, SOX2 (transcription factor for stemness) and cyclin E1, Cyclin A1, cyclin B1 (cell cycle cyclins) and E2F1, mTOR, PCNA proliferative marker gene expression were decreased following the knockdown of CSNK2β (Figure 4B). These results showed the critical role of CSNK2β in the expression of genes associated with cell cycle, cancer stemness, and metastasis.

**Figure 4.**
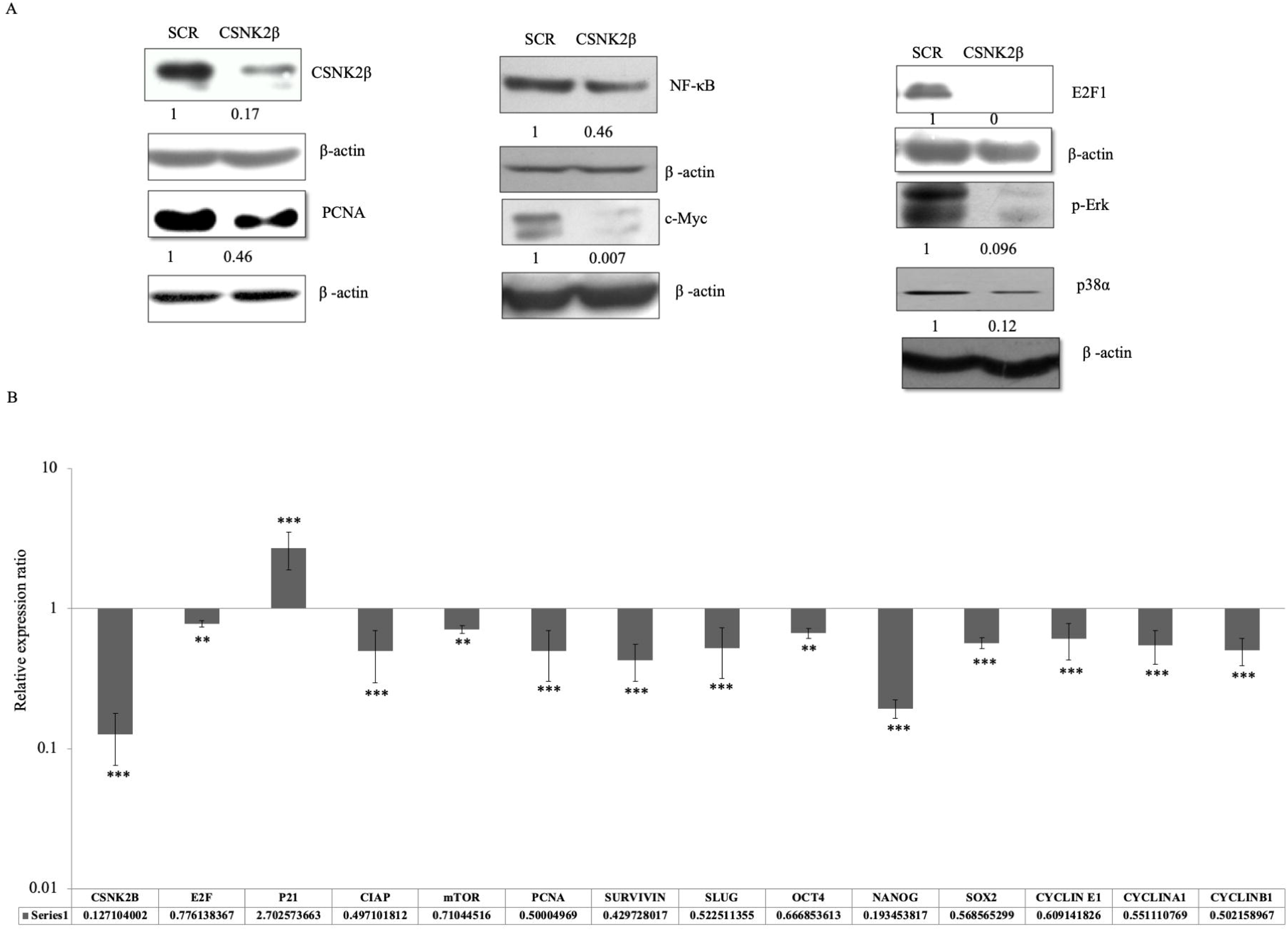
Silencing of CSNK2β modulated key markers associated with cell proliferation, migration and cell stemness. **(A)** Western blot analysis of proteins associated with cell proliferation. (B) qPCR analysis of genes associated with apoptosis, cell cycle, cell stemness and metastasis. Data are represented as mean ± SD from triplicate samples. Data are represented as mean ± SD from triplicate samples, where **p <0.01 and ***p < 0.001

## Discussion

Breast cancer is the most commonly diagnosed as well as leading cause of cancer deaths in women^24^. Mortality due to breast cancer has been decreasing due to early diagnosis, improved adjuvant therapy and low usage of hormone replacement therapy^25^. Despite all these advancements and outcomes triple negative breast cancer (TNBC) characterised by absence of molecular markers-estrogen receptor, progesterone receptor and epidermal growth factor 2, still remains a major challenge. Statistics show that TNBC constitute 15% of all breast cancer^26^ and triple negative breast cancer patients have a decrease in survival after initial 3-5 years of diagnosis^27^. Thus it is indispensable to explore the potential molecular target which is critically involved in maintaining cancer phenotype and significantly contribute to metastasis, morbidity, and mortality due to breast cancer. In our study, we chose to elucidate the role of CSNK2β in breast cancer (MDA-MB-231, a TNBC cell line) in *in-vitro*.

Our results showed that knockdown of CSNK2β significantly decreased the cell viability and colony growth of MDA-MB-231 cells. On the basis of these observations, we next evaluated the status of ROS inside the MDA-MB-231 cells since controlled production of ROS is beneficial for proliferation and viability of cancer cells but a disproportional increase of ROS can induce cell cycle arrest and at its peak can induce apoptosis of cancer cells^28,29^. In our study we found that knockdown of CSNK2β markedly induced ROS production which seems to be consistent with earlier that use of a pharmacological inhibitor which blocks catalytic activity of CK2 induced apoptosis in leukemic cell lines via up regulation of intracellular H2O2^30^. Moreover we found the G2M cell cycle arrest of MDA-MB-231 cells following knockdown of CSNK2β and in some studies it has been shown that ROS accumulation is at its peak in the G2M phase of cell cycle^31^. So accumulated ROS found in this study can be attributed to arrest of cells at G2/M phase following knockdown of CSNK2β. However the exact mechanism which triggered redox modulation upon knockdown of CSNK2β was not explored in this study and further investigation is needed to understand the association of CSNK2β and the ROS status inside MDA-MB-231 cancer cells.

A perturbed cell cycle progression may lead to apoptosis^32^. In our study, we found siRNA knockdown of CSNK2β in MDA-MB-231 led to nuclear condensation and change of cellular morphology, a characteristic of cell undergoing apoptosis. Further this study also showed that depletion of CSNK2β modulated molecular markers associated with apoptosis and autophagy, which is consistent with earlier reports of involvement of CK2 in suppressing apoptosis, as over-expression of CK2 in prostate cancer cells prior to treatment with etoposide rescued against cell death^33^ and down regulation of CK2 (α subunit) leads to autophagic cell death in glioblastoma cell lines^34^. These results suggest that CSNK2β has a prominent role in survival of MDA-MB-231 cells by suppressing the apoptotic and autophagy machinery.

In this study we also explored the role of CSNK2β in the metastasis and invasion, as these two are important phenotypes associated with breast cancer. Metastasis, which is evident from stage 2 onwards and nearly 30% women develop metastasis even after initial diagnosis^35^, requires identification of markers involved in this lethal property of cancer. In present study we found that silencing of CSNK2β markedly up-regulated E-cadherin as loss of later cause invasiveness in carcinomas and other cancers^36^. In search of other markers, associated with migration and hence metastasis, being affected following inhibition of CSNK2β we found down regulation of β1 Integrin, MMP-9, Vimentin and β-catenin. β1 Integrin, was found to be inhibited in CSNK2β knockdown cells which is a majorly expressed integrin and has been reported to actively involved in metastasis and has attracted a considerable attention as a target for immunotherapy in breast cancer^37^. MMP-9, a protease, facilitate cancer cells in degrading extra cellular matrix for metastasis and invasion, has been found to be over expressed in many cancers including breast cancer. In this study we found MMP-9 to be inhibited following knockdown of CSNK2β which corroborates with previous finding of MMP-9 inhibition with chemically inhibited CK2 in lung adenocarcinoma cells^38^. Vimentin, a major factor associated with epithelial mesenchymal transition (EMT)^39^ and a vital prognostic biomarker and therapeutic target correlated with EMT in TNBC^40^ was found to be down regulated in CSNK2β knockdown cell. In addition to these we also found that knockdown of CSNK2β down-regulates the expression of β-.catenin It has been previously shown that β-.catenin plays pivotal role in tumor related phenotypes including migration and stemness in particular in TNBC cells^41^. Collectively these results showed that CSNK2β mediates migration and invasion of MDA-MB-231cells and these critical markers appear to be acting downstream of CSNK2β.

To get the further insight into the biological significance of CSNK2β in pathogenesis of breast cancer, we performed the expression study of the key molecules related to cell proliferation, survival, cell cycle, apoptosis and autophagy-related genes by western blotting and real time PCR.

Raf/MEK/ERK pathways are activated in many tumors (prostate, breast, leukemia, melanoma, thyroid) which transmit the signals from cell surface receptors to transcription factors and can be exploited for therapeutic intervention^42^. In the present study, we found that CSNK2β regulates the expression of p-ERK, p38-α, c-MYC proteins which are connected to MAPK pathway, in MDA-MB-231 cells. These data suggested that targeting CSNK2β might be a potential strategy to improve clinical outcomes in future. PCNA has been earlier reported as a reliable marker to access the growth and predicting the prognosis in breast cancer^43^. Thus we investigated the effect of silencing of CSNK2β on PCNA and found its downregulation at both transcript and protein level. A recent study on the interbreeding of MMTV-PyMT mice with E2F1, E2F2, or E2F3 knockout mice showed that in addition to cell cycle control E2F targets a number of genes related to angiogenesis, extracellular matrix modification, proliferation and survival of tumor cells which was important for metastasis^44^. We also found that CSNK2β reduces the expression of E2F1 which might affect a large fraction of genes in breast cancer. NF-κB expression leads to the induction of genes related to apoptosis, cell cycle, cell invasion which contributes to tumorigenesis, chemoresistance and radioresistance^45^. Our result showed that CSNK2β might regulate the cell proliferation through the NF-κB pathway.

One of the most striking results obtained in this study following silencing of CSNK2β was a highly significant up regulation of p21 at transcript level. p21 is well known as sensor and effector of multiple growth inhibiting signals and its over expression has been reported to cause arrest in breast carcinoma cell lines^46^. This result is in similar line of earlier report where inhibition of CK2 using inhibitor caused up-regulation of p21 in glioblastoma cells and induced G2M arrest of cell cycle^47^. Knockdown of CSNK2β also led to down-regulation of mTOR. Alteration in mTOR pathway is a common anomaly in various breast cancer subtypes^48^. We also checked the expression of survivin and cIAP as both are inhibitors of apoptosis and overexpressed in majority of breast cancer and survivin has been found to confer resistance to chemotherapy^49^.

In this study we also found that expressions of slug, oct4, nanog and sox2 were significantly down regulated at transcript level. Slug over expression has been reported to induces stemness and promote migration in hepatocellular carcinoma, breast and other cancer types^50,51^. Slug overexpression is associated with reduced expression of E-cadherin,lymph node metastasis and poor survival of patients^52^. Interestingly, in our study we found that knockdown of CSNK2β led to downregulation of slug and upregulation of E-cadherin. Sox2, a transcription factor, plays important role in pluripotency of stem cells. Down regulation of Sox2 is a promising outcome of this study as Sox2 has been reported to positively influence cell proliferation and migration of TNBC cells^53^. Down regulation of oct-4 and nanog is another crucial finding of this study, since both of the pluripotency factors are highly expressed in multiple cancers^54,55^ including various subtypes of breast cancer and their abrogation reverses epithelial mesenchymal transition in lung carcinoma^55^. Although this is very initial finding for relationship between CSNK2β and mentioned stemness marker, a further study is needed to explore the signal relationship among them.

Present study also explored the expression level of cyclins (A1, B1 and E1) at transcript level-, following knockdown of CSNK2β. These cyclins have been shown to be associated with breast cancer progression, via modulating different cellular pathway such as by modulating VEGF and hence angiogenesis^56^ and poor survival of patients ^57,58^.

In conclusion the present study explored the participation of β subunit of CK2 in breast cancer pathogenesis using TNBC cell line MDA-MB-231, since most of the studies have been carried out only on α subunit. Our study has shown that silencing of CSNK2β significantly inhibited growth of MDA-MB-231 cells, caused cell cycle arrest and induced apoptosis. Further this study also found that CSNK2β knockdown significantly hampered migration and invasion of MDA-MB-231 cells via down regulation of MMP-9, vimentin, beta-integrin 1 and up regulation of E-cadherin. We found that β subunit of CK2 participate in contributing cancer pathogenesis in a similar manner as α subunit of CK2 in some aspect of cancer progression as mentioned in the text.

## Author Contribution

Shibendra Kumar Lal Karna (SKLK) executed most of the experiments and written manuscript, Bilal Ahmad Lone (BAL), performed some experiments and manuscript writing, Faiz Ahmad (FA) contributed some experiments and manuscript writing, Nerina Shahi (NS) performed part of the experiments, Yuba Raj Pokharel (YRP) designed, supervised and edited manuscript.

